# Neuroinvasion by Simian Immunodeficiency Virus Triggers Glial Senescence and Accelerated Neurodegeneration

**DOI:** 10.64898/2025.12.15.694426

**Authors:** Miranda D. Horn, Cecily C. Midkiff, Alison R. Van Zandt, Ahmad A. Saied, Andrew G. MacLean

## Abstract

Virus-induced accelerated aging has emerged as a potential contributor to HIV-associated neurocognitive disorders (HAND), despite widespread implementation of combination antiretroviral therapy (cART). Although evidence of accelerated aging in people living with HIV (PLWH) has been reported, most investigations of acute infection rely on in vitro systems or small animal models, leaving a critical gap in understanding early neuropathological events. To address this, we analyzed formalin-fixed, paraffin-embedded (FFPE) brain tissues from rhesus macaques acutely infected with simian immunodeficiency virus (SIV). We focused on two key aging-related proteins: the cellular senescence marker p16^INK4a^ (p16) and the NAD-dependent deacetylase sirtuin 1 (SIRT1). We hypothesized that accelerated aging phenotypes would be detectable during acute infection, manifesting as increased p16 expression and altered SIRT1 levels, correlating with neurodegeneration. Consistent with this hypothesis, we observed marked upregulation of GFAP and p16, along with evidence of neurodegeneration, across multiple brain regions - including the frontal lobe, caudate, putamen, thalamus, hippocampus, and cerebellum - by 21 days post-infection. These findings suggest that aging-related and senescence pathways are activated almost immediately following HIV infection, highlighting the potential importance of astrocyte- or CNS-specific therapeutic strategies to mitigate early neuropathology.

## Introduction

Access to combination antiretroviral therapy (cART) has greatly improved the lifespan of people with HIV (PWH) and reduced the incidence of HIV-associated dementia. Despite effective viral suppression, the prevalence of HIV-associated brain injury (HABI) has remained high, even in individuals with successful viral suppression in the periphery^1^. PWH also display accelerated rates of biological aging in the brain, as indicated by brain imaging and cognitive assessments^2–4^ as well as biological markers of aging and neurodegeneration^5,6^. These findings suggest that HIV infection has long-lasting effects on the CNS that are not fully mitigated by cART. The underlying mechanisms driving these alterations in brain aging remain an area of active debate. Several hypotheses have been proposed, including (1) damage from early CNS insult during acute infection, (2) ongoing damage due to persistent viral reservoirs, (3) incomplete suppression of CNS inflammation by cART, and (4) potential neurotoxicity associated with long-term cART use^7^. Clarifying the relative contributions of these factors is critical to developing interventions that can preserve cognitive function and brain health in PWH. Thus, models that allow us to examine the early events in HIV infection are required.

Similar patterns of altered brain aging have been reported in Simian Immunodeficiency Virus (SIV)-infected rhesus macaques (RMs), an established model for studying HIV neuropathogenesis^8,9^. Studies in SIV-infected RMs have demonstrated accelerated or atypical aging trajectories in the brain, marked by structural and functional changes^2,10,11^, behavioral impairments^12–17^, and molecular signatures consistent with neurodegeneration^18,19^. Our laboratory has recently demonstrated that while cART successfully reduces neurodegeneration relative to chronically SIV-infected RMs in the frontal lobe, it has a limited effect on markers of aging or neurodegeneration in other brain regions^20^.

In this study, we focus on the hypothesis that early CNS injury – occurring during acute SIV infection and before cART would normally be initiated – is a primary driver of altered brain aging. To test this, we assess biological markers of aging and neurodegeneration in acutely infected animals compared to naïve controls. By isolating the effects of early infection in the absence of long-term cART exposure or chronic inflammation, this work aims to clarify the timing and origin of neurobiological changes that contribute to premature or atypical brain aging in the context of HIV/SIV.

## Materials and Methods

### Ethics Statement

Animals were handled in accordance with the American Association for Accreditation of Laboratory Animal Care and NIH “Principles of laboratory animal care”. Veterinarians and their staff carried out all animal procedures as approved by the Institutional Animal Care and Use Committee of Tulane University. Animals were humanely euthanized by veterinary staff by anesthesia with ketamine hydrochloride (10 mg/kg) followed by an overdose with sodium pentobarbital and necropsy, in accordance with established endpoint policies. Tissues were fixed for 48 hours in 10% neutral buffered formalin with zinc modification followed by paraffin embedding.

### Selection of Animals and Tissues

Formalin-fixed paraffin-embedded (FFPE) tissues from six RMs, aged 3.94 to 4.81 years of age were used in this study. Three of these animals were naïve to SIV and cART and were healthy at the time of investigator-initiated euthanasia (Naïve) and the other three animals had been infected with SIVmac239 for 21-22 days prior to investigator-initiated euthanasia (Acute SIV).

The frontal lobe, hippocampus, caudate, putamen, thalamus, and cerebellum have previously been identified as brain regions that show atrophy in PLWH and SIV-infected macaques^2,10,21–23^. Thus, we used H&E slides to select FFPE tissue blocks that contained these brain regions for each animal whenever possible. (see Table 1 for more detailed information on the animals and specific brain regions analyzed for each animal).

**Table 1:**
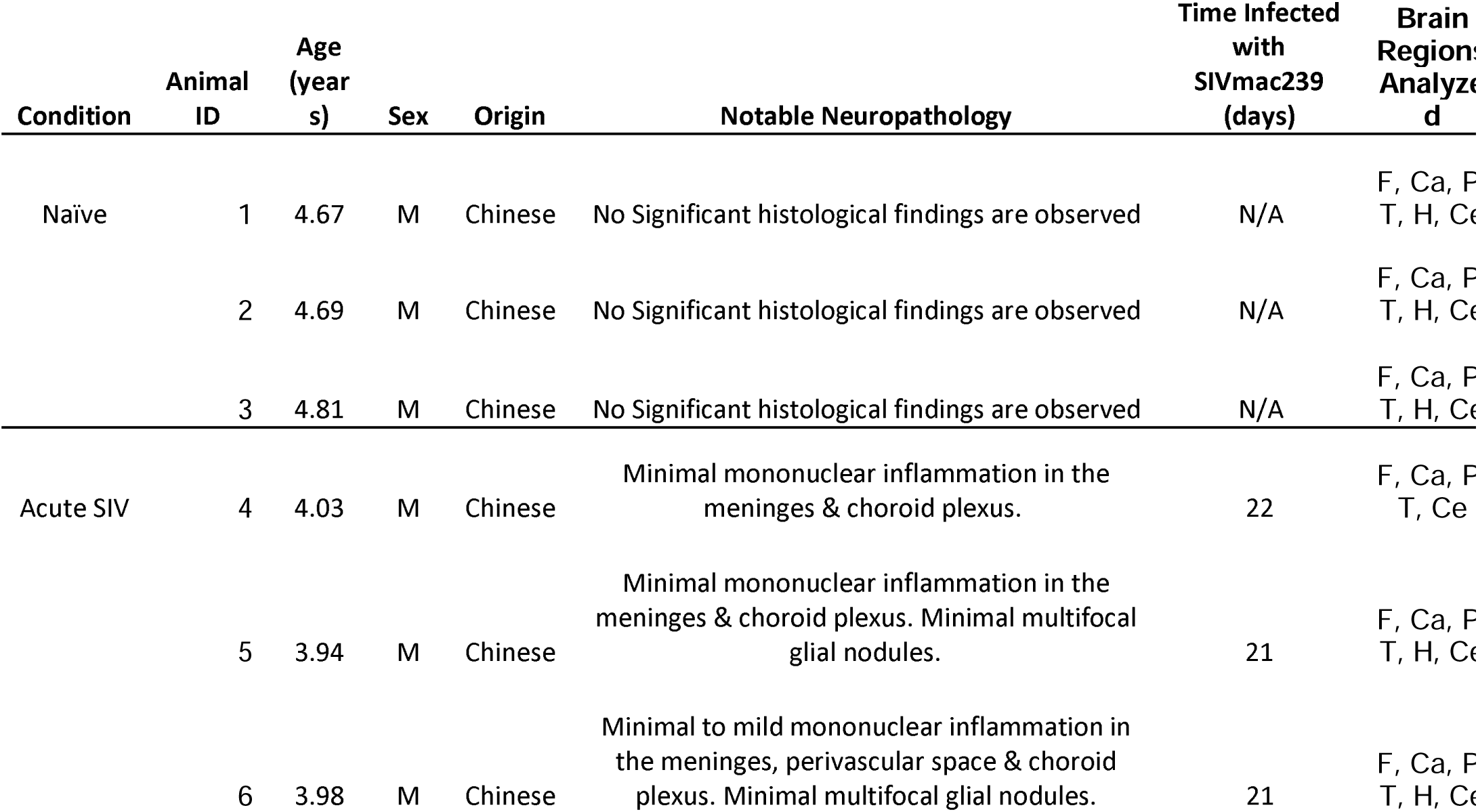
Information on the animals used in this study. Abbreviations: F = Frontal lobe, Ca = Caudate, P = Putamen, T = Thalamus, H = Hippocampus, Ce = Cerebellum.

| Condition | Animal ID | Age (years) | Sex | Origin | Notable Neuropathology | Time Infected with SIVmac239 (days) | Brain Regions Analyzed |
| --- | --- | --- | --- | --- | --- | --- | --- |
| Naïve | 1 | 4.67 | M | Chinese | No Significant histological findings are observed | N/A | F, Ca, P<br>T, H, Ce |
|  | 2 | 4.69 | M | Chinese | No Significant histological findings are observed | N/A | F, Ca, P<br>T, H, Ce |
|  | 3 | 4.81 | M | Chinese | No Significant histological findings are observed | N/A | F, Ca, P<br>T, H, Ce |
| Acute SIV | 4 | 4.03 | M | Chinese | Minimal mononuclear inflammation in the meninges & choroid plexus. | 22 | F, Ca, P<br>T, Ce |
|  | 5 | 3.94 | M | Chinese | Minimal mononuclear inflammation in the meninges & choroid plexus. Minimal multifocal glial nodules. | 21 | F, Ca, P<br>T, H, Ce |
|  | 6 | 3.98 | M | Chinese | Minimal to mild mononuclear inflammation in the meninges, perivascular space & choroid plexus. Minimal multifocal glial nodules. | 21 | F, Ca, P<br>T, H, Ce |

### Immunohistochemistry

The TNPRC Confocal Microscopy and Molecular Pathology Core performed chromogenic immunohistochemistry for Iba1 for analysis of microglia/macrophages as previously described^20,24^. Briefly, slides containing 5µm-thick tissues were baked overnight at 60°C prior to being deparaffinized and rehydrated. Antigen retrieval was performed using a Tris-based solution, pH 9 (Vector Labs H-3301) with 0.1% Tween20, deionized water, and a citrate-based solution, pH 6.0 (Vector Labs H-3300). Slides were washed in phosphate-buffered saline, deionized water, and Roche reaction buffer prior to being loaded onto the Ventana Discovery Ultra autostainer for blocking, primary antibody (rabbit anti-IBA, Wako 019-19741, 1:3000 dilution) incubation, washing, secondary antibody incubation (OMap anti-Rb HRP, Ventana, cat. #760-4311), washing, DAB color development, and counterstaining with hematoxylin II. Upon removal, slides were washed with deionized water containing 0.1% Dawn dish soap and plain deionized water for a total of 5 cycles. Slides were then cleared before being permanently mounted with StatLab™ AcryMount Plus Mounting Media (FisherScientific, cat. # STSL80PLUS4) and left overnight to dry prior to imaging.

### Immunofluorescent Staining

For all immunofluorescent staining, slides were stained as previously described ^20,24^. Briefly, slides containing 5 µm-thick FFPE tissues were baked at 60°C overnight and deparaffinized prior to antigen retrieval (Antigen Unmasking Solution, Citric Acid Based, Vector Laboratories, Inc., cat. #H-3300), blocking [normal goat (MP Biomedicals, LLC., cat. #2939149) or normal donkey (GeminiBio, cat. #100-151) serum], and antibody application (see Table 2 for antibody information). Slides were washed in PBS-FSG-Tx100 (1X Phosphate-Buffered Solution diluted from 10X Phosphate-Buffered Solution, Fisher BioReagents™, cat. #BP 3994; 0.2% Fish-Skin Gelatin, Sigma Aldrich, cat. #1002923460; 0.1% Triton-100, Fisher BioReagents™, cat. #BP 151 500) and 1X PBS-FSG after each antibody, and with 1X PBS prior to applying mounting media (EverBrite TrueBlack® Hardset Mounting Medium, Biotium, cat. #23018) and coverslips.

**Table 2:**
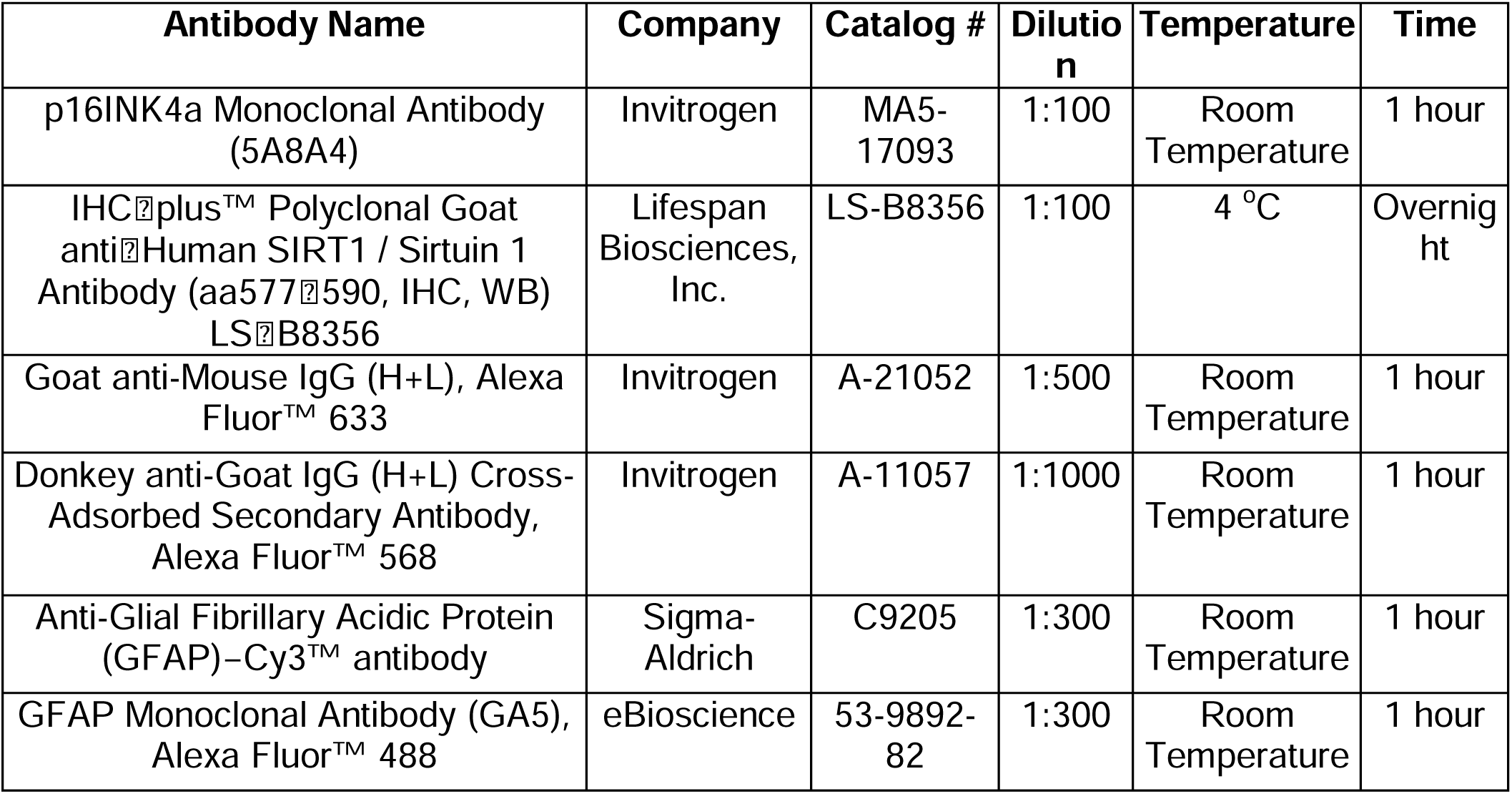
Information on the antibodies used for immunofluorescent staining.

### FluoroJade C staining

FJC (Fluoro-Jade C Staining Kit with DAPI Counter Stain, Histo-Chem Inc., cat. #FJC-SK-DAPI) staining was completed as previously described^20,24^. Briefly, following deparaffinization, slides were incubated in 70% alcohol and deionized water prior to dampening of autofluorescence by potassium permanganate. Slides were then washed in deionized water and stained with FJC and DAPI. Slides were again washed in deionized water before being dried in the oven at 60°C and cleared in xylene prior to application of mounting media (Micromount mounting media, Leica Biosystems, cat. #3801730) and coverslips. Slides were allowed to set at room temperature overnight before imaging.

### Image Analysis

A Zeiss AxioScan.Z1 slide scanner at 20x magnification was used for GFAP, p16, and SIRT1 imaging. As previously described^20,24^, all slides for a given brain region and stain were scanned using the same settings. HALO® Image Analysis software (Indica Labs) was used for analysis of whole-slide images by annotating appropriate regions of interest and setting parameters in the Highplex FL module (Indica Labs) to identify phenotypes of interest. Summary data were exported to Microsoft Excel and used to determine the percentage of marker-positive cells, dual-labeled cells, and relative fluorescent intensity for each marker. A Hamamatsu NanoZoomer360 at 40x magnification was used for Iba1 imaging. The Microglia Activation Module in HALO® Image Analysis was used to determine the percentage of Iba1+ cells and activated Iba1+ cells.

### Statistical Analyses

To assess the differences in marker expression between groups, the percentage of marker-positive cells, co-labeled cells, and relative intensities were compared using unpaired t-tests in Prism GraphPad (version 10, GraphPad Software, La Jolla, CA). To determine the relationship between expression of multiple markers, two-tailed Pearson’s correlation coefficients were calculated using Prism GraphPad. A p value < 0.05 was considered significant for all analyses.

## Results

### Description of pathology from selected animals

Animals infected with SIV develop perivascular inflammation, meningitis, and microglial nodules in the brain. Detailed examination revealed minimal inflammation in the meninges and choroid plexus of acutely infected animals (Table 1). While there were increased numbers of Iba1+ cells in all brain regions except for thalamus, when whole sections were imaged, none of the differences in the percentage of Iba1+ cells was significant (Figure 1A).

**Figure 1.**
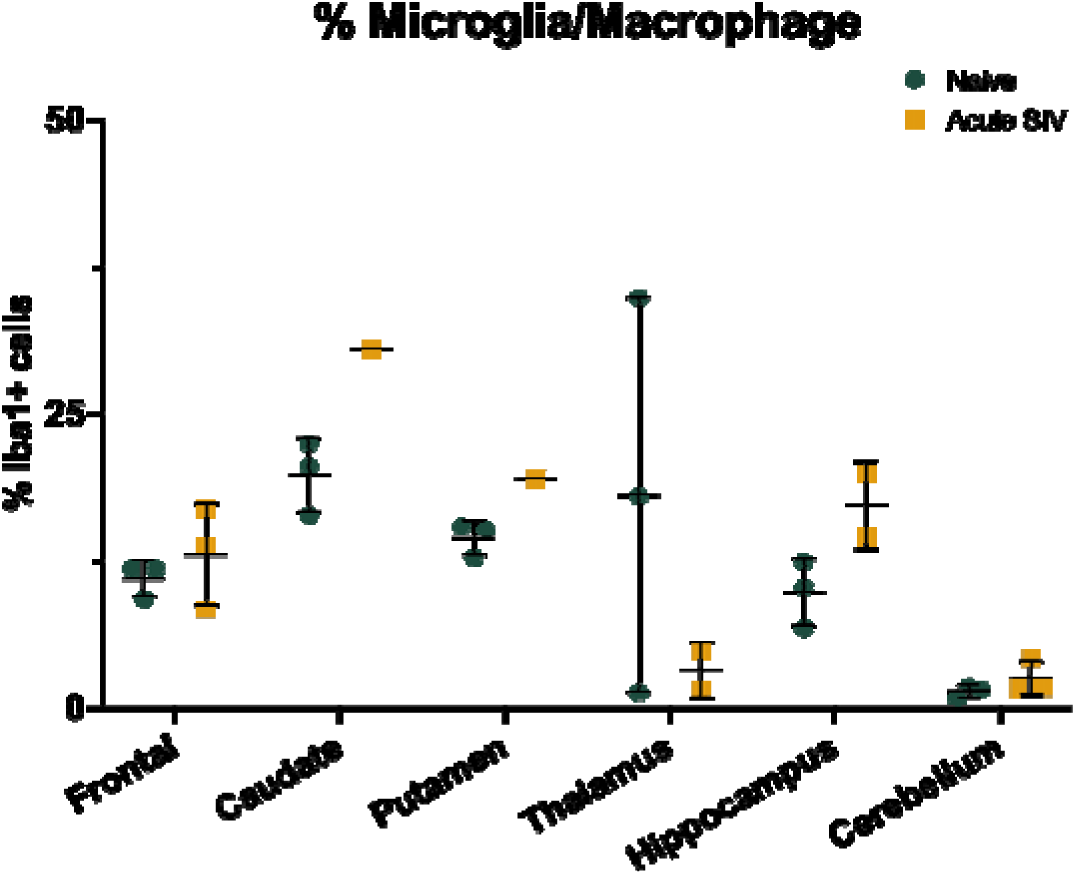
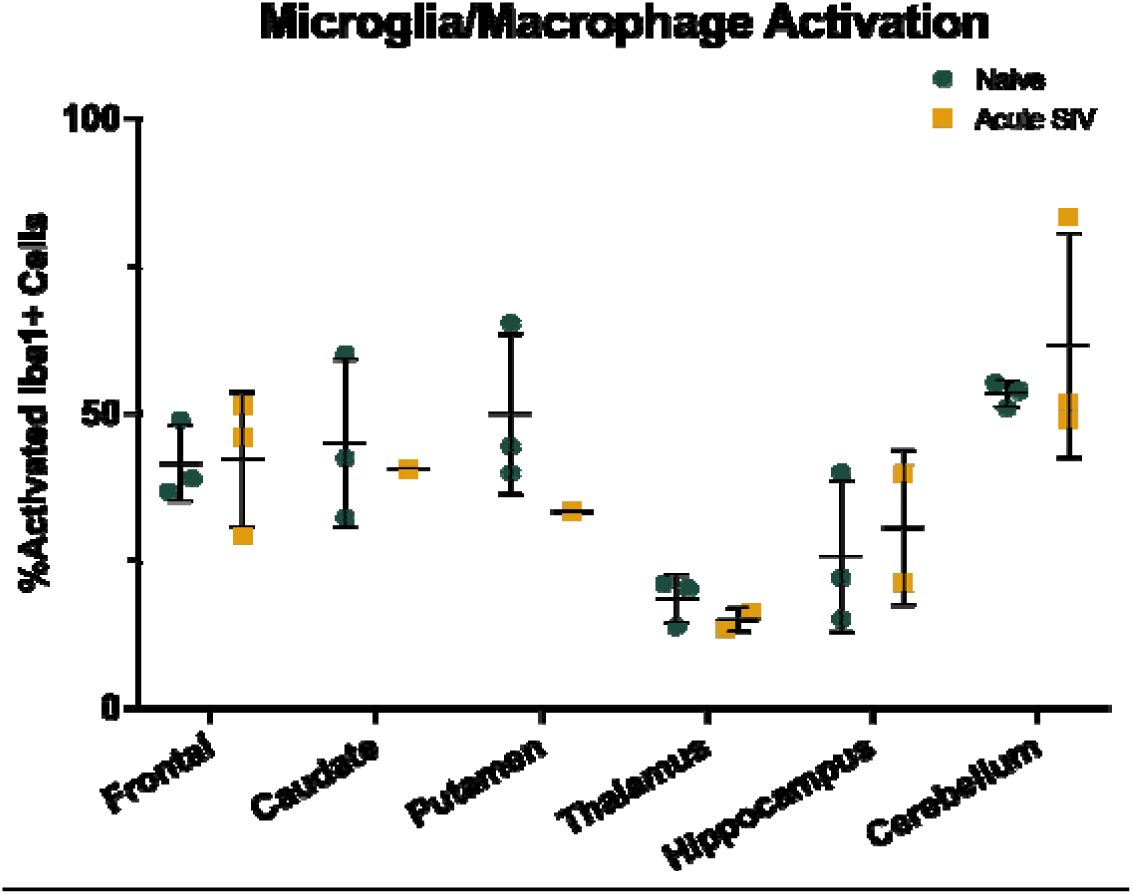
Microglia activation is unaltered in acute SIV infection. (A) The percentage of Iba1+ cells does not change in acute SIV infection across all observed brain regions. (B) The percentage of activated Iba1+ cells does not change with acute SIV across all observed brain regions. All data points are presented and mean +/− SD are plotted for each group. Each brain region was analyzed independently using unpaired t-tests.

Similarly, no significant differences in the percentage of activated microglia within specific brain regions (Figure 1B) was noted.

### GFAP expression increases with acute SIV infection across the brain

As astrocytes can be activated during acute infections, we expected to see increases in GFAP expression in our acutely infected animals relative to uninfected animals. To test this hypothesis, we assessed changes in the percentage of GFAP+ cells and the intensity of GFAP across six brain regions (Figure 2). We found that the percentage of GFAP+ cells was indeed increased during acute SIV infection in the caudate, putamen, hippocampus, and cerebellum (Figure 2A), but not in frontal lobe nor thalamus.

**Figure 2.**
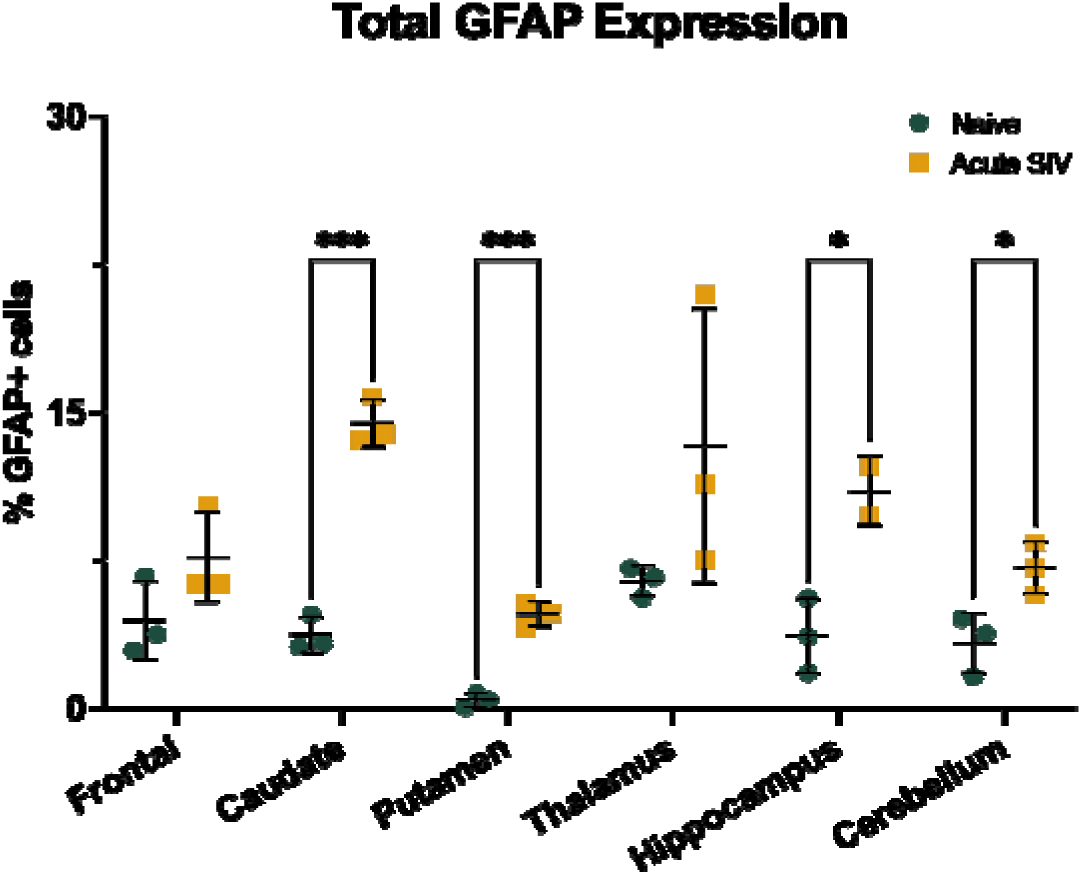
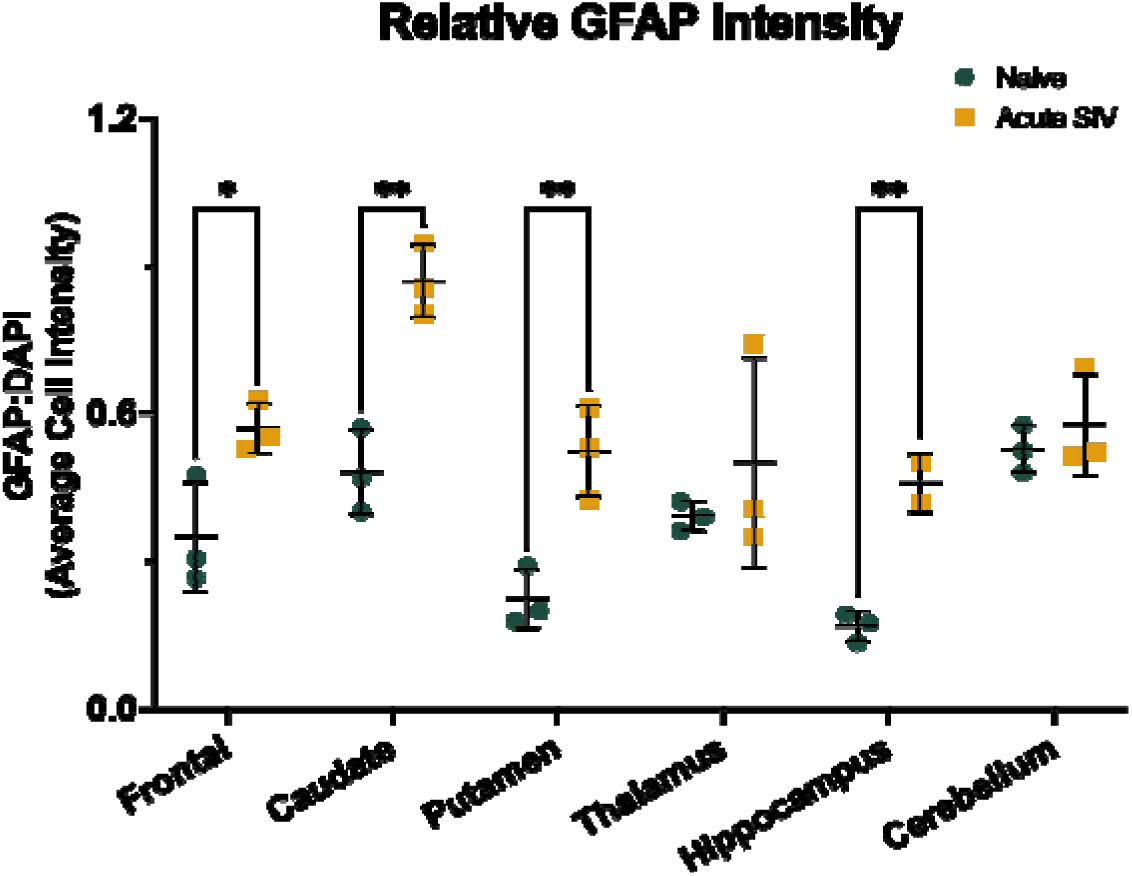
Acute SIV infection increases GFAP expression across the brain in rhesus macaques. (A) Acute SIV infection increased the percentage of GFAP-positive cells in the caudate, putamen, hippocampus, and cerebellum. (B) Relative GFAP intensity was increased in the frontal lobe, caudate, putamen, and hippocampus of acute-SIV infected animals. All data points are presented and mean +/− SD are plotted for each group. Each brain region was analyzed independently using unpaired t-tests. * p < 0.05, ** p <0.01, *** p < 0.001.

As we have previously found GFAP intensity to better correlate with aging in the RM^24^, we hypothesized that if SIV-induced biological aging begins during acute infection then GFAP intensity would also increase in these animals. Here we found GFAP intensity to increase in acute SIV infection in frontal lobe, caudate, putamen, and hippocampus (Figure 2B).

While this increase in GFAP may be due to early inflammatory responses, if left unresolved it could contribute to SIV-induced biological aging.

### Biological aging marker p16INK4a is increased across multiple brain regions during acute SIV infection

To assess the effect of acute SIV infection on biological aging, we first measured the expression of the biological aging marker p16 throughout the brain. We have previously found p16 to increase with age^24^ and chronic SIV infection^20^, indicative of SIV-induced biological aging that was not prevented by cART, thus we hypothesized that this may be triggered during acute infection, prior to initiation of cART. Here, we observed increases in p16 expression during acute SIV infection across the brain (Figure 3). The percentage of p16+ cells was significantly increased in the caudate, putamen, thalamus hippocampus, and cerebellum during acute infection relative to naïve animals (Figure 3A).

**Figure 3.**
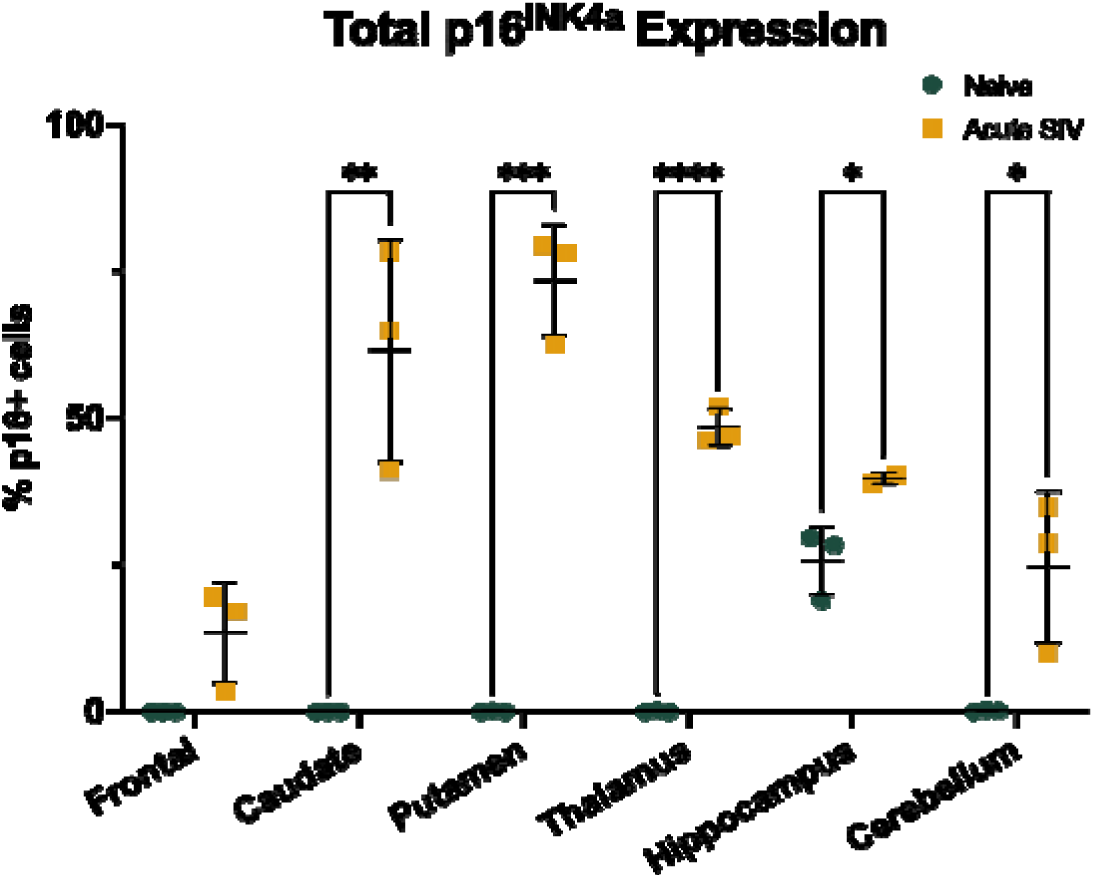
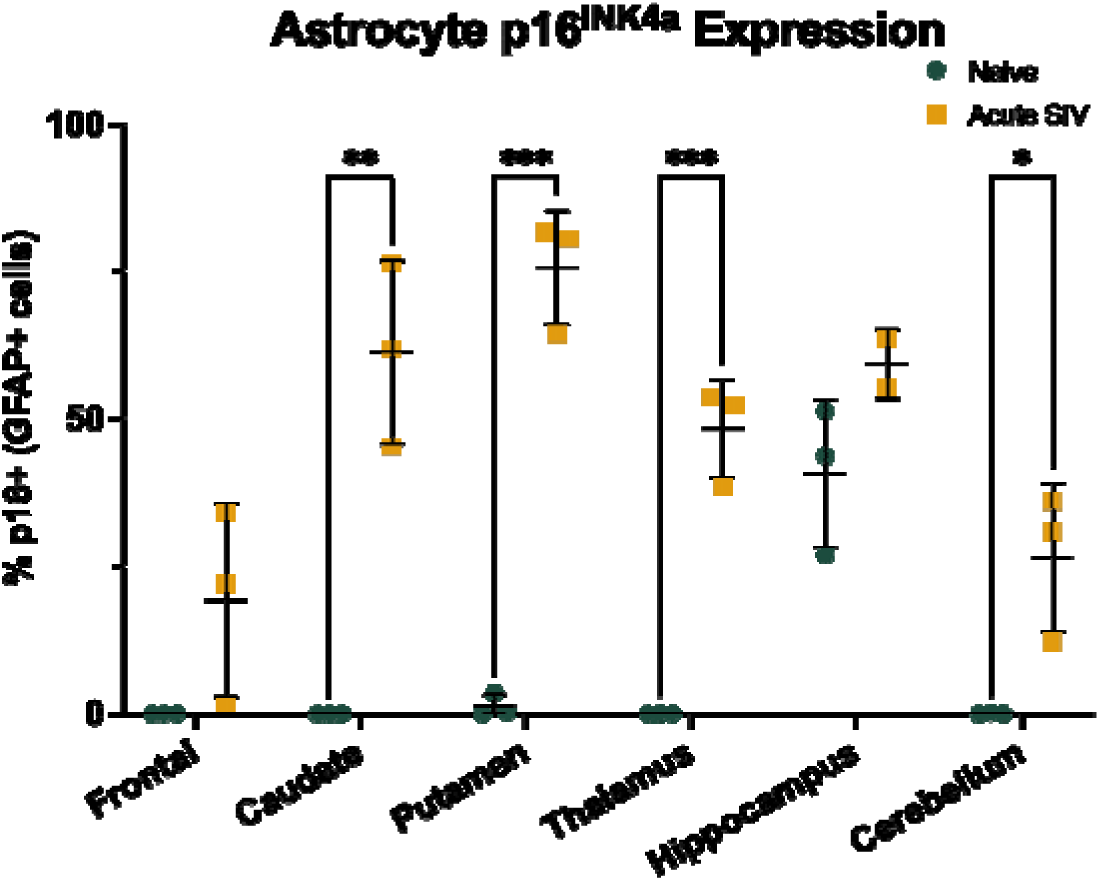
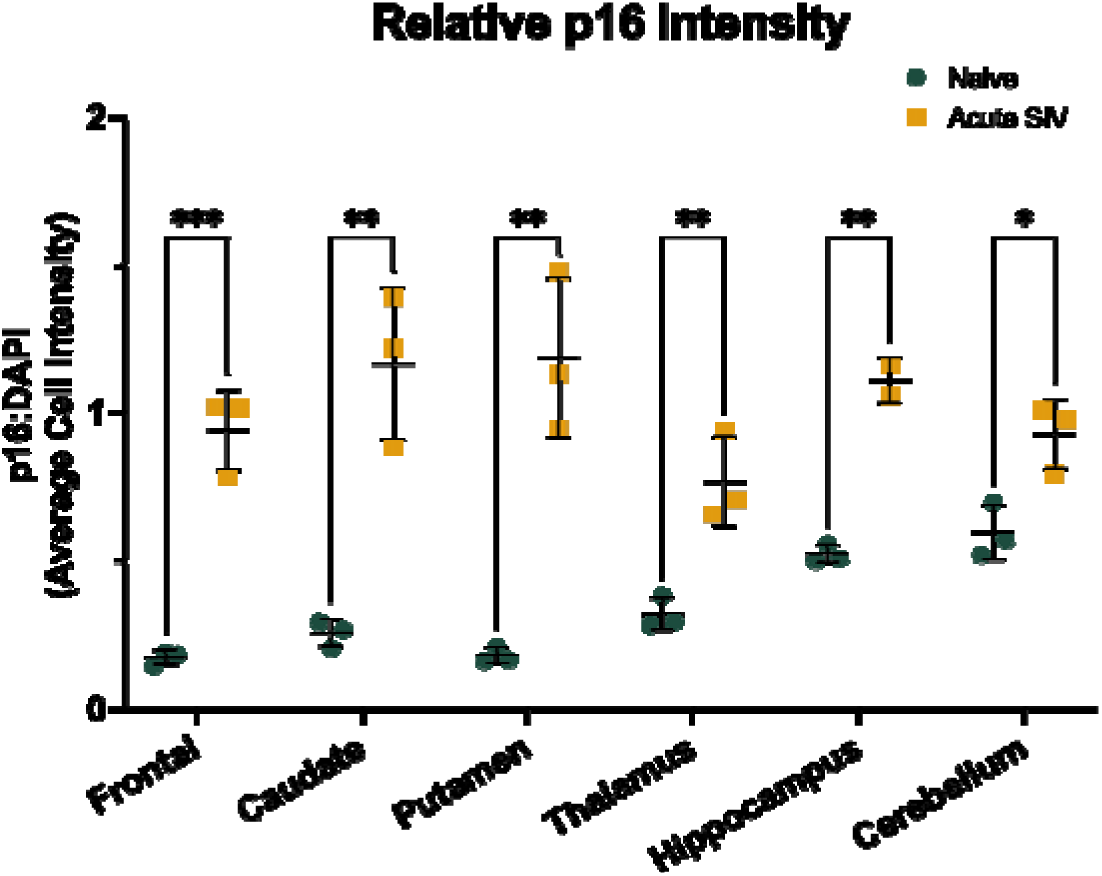
Acute infection with SIV induces increased p16^INK4a^ expression across the brain of rhesus macaques. (A) Acute infection with SIV increases p16 expression in the caudate, putamen, thalamus, hippocampus, and cerebellum. (B) Astrocytes show a similar pattern, with increased p16 expression in the caudate, putamen, thalamus, and cerebellum. (C) The relative intensity of p16 staining increased across all brain regions following acute SIV infection. All data points are presented and mean +/− SD are plotted for each group. Each brain region was analyzed independently using unpaired t-tests. * p < 0.05, ** p <0.01, *** p < 0.001, ****p < 0.0001.

We saw a similar pattern in GFAP+ cells (Figure 3B), with increases in the percentage of p16+ astrocytes in the caudate, putamen, thalamus, and cerebellum of acutely infected animals.

Finally, we also observed an increase in the intensity of p16 staining during acute SIV infection (Figure 3C) in all six brain regions examined.

### SIRT1 expression decreases in the hippocampus following acute SIV infection

SIRT1 expression positively correlates with neurodegeneration in the frontal lobe of uninfected animals^24^, and in the cerebellum of chronically-infected RMs^20^. In contrast, we saw a significant decrease in the percentage of SIRT1+ cells relative to naïve animals in the hippocampus of acutely-infected RMs (Figure 4A), with no significant difference in any other region examined.

**Figure 4.**
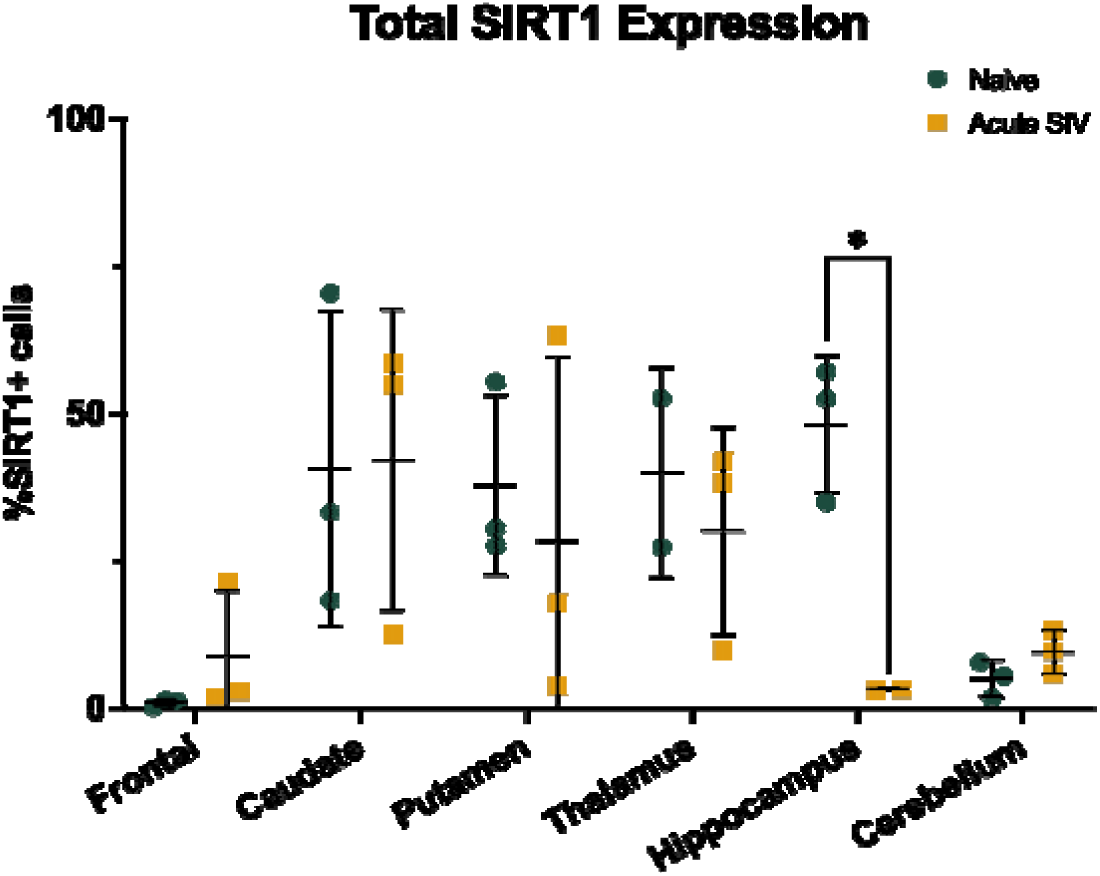
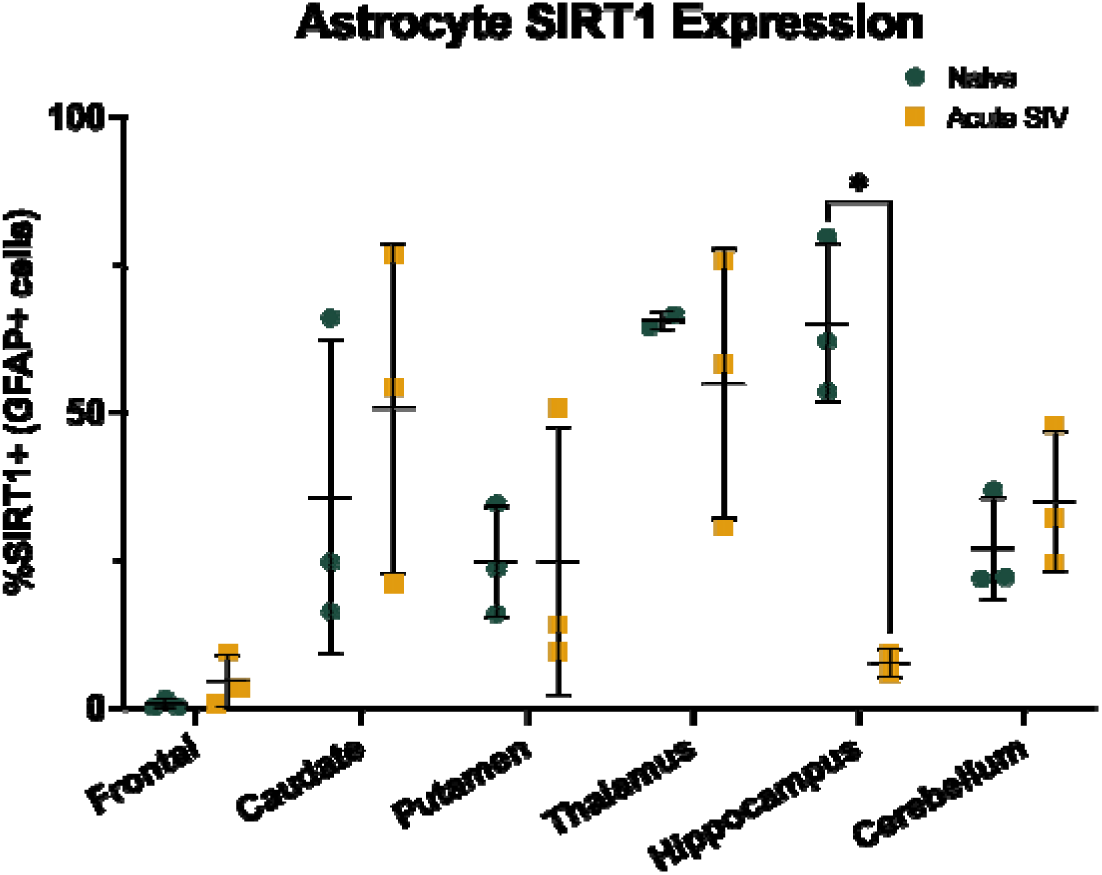
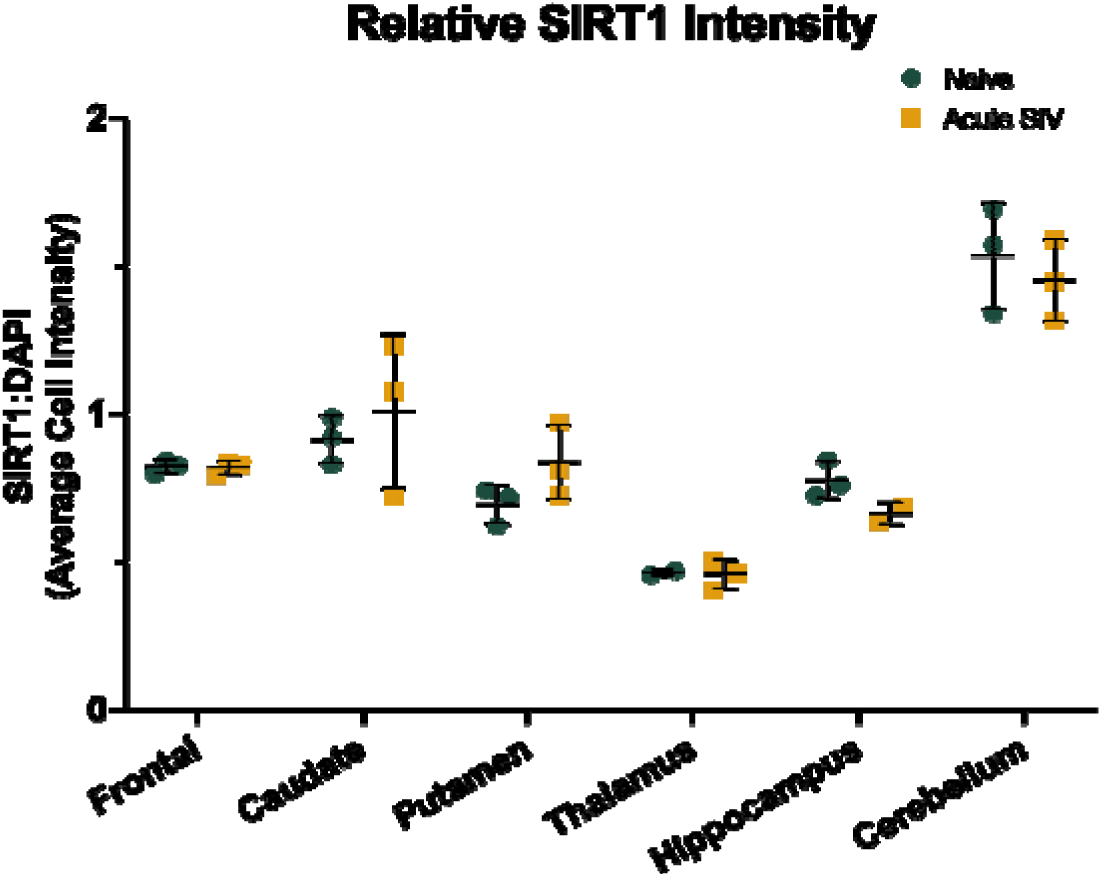
Acute infection with SIV reduces SIRT1 expression in the hippocampus of rhesus macaques. (A) The percentage of SIRT1+ cells decreased in the hippocampus following acute SIV infection, no other differences were noted. (B) Astrocytes in the hippocampus similarly had reduced SIRT1 expression following acute SIV infection (C) The relative intensity of SIRT1 was unaltered with acute SIV infection across all brain regions. All data points are presented and mean +/− SD are plotted for each group. Each brain region was analyzed independently using unpaired t-tests. * p < 0.05.

This same pattern was observed in astrocytes (Figure 4B)

but there were no significant changes in the intensity of SIRT1 staining relative to DAPI staining in any regions examined (Figure 4C).

### FluoroJade C staining is increased in acutely SIV-infected animals in a region-specific manner and correlates with biological aging markers

Neurodegeneration was increased in the putamen, thalamus, and hippocampus, as measured by the relative intensity of FJC staining (p = 0.0461, p = 0.0043, and p = 0.0002, respectively), but not the percentage of FJC+ cells (Figure 5).

**Figure 5.**
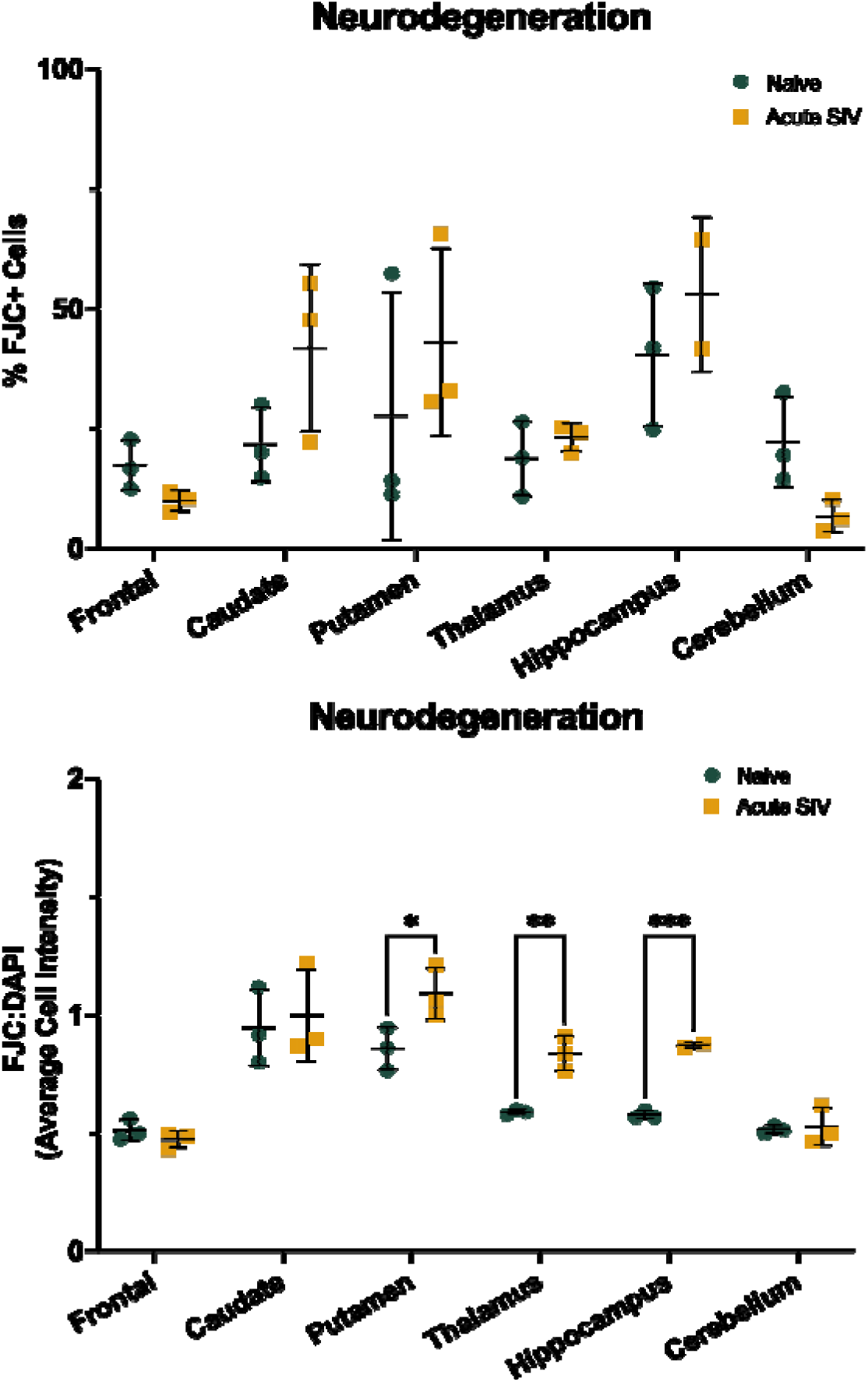
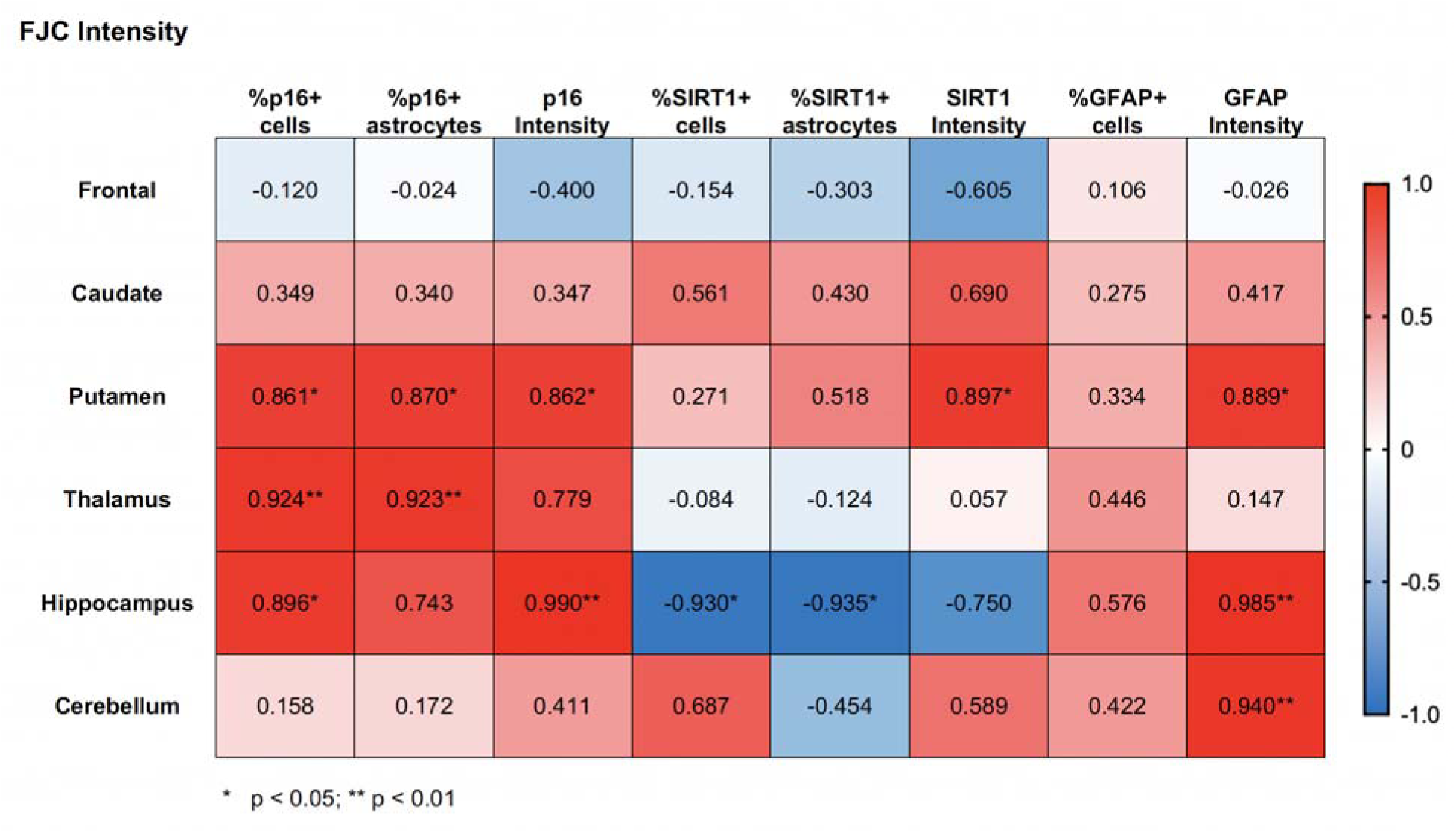
Neurodegeneration occurs after acute SIV infection in multiple brain regions and correlates with markers of biological aging. (A) Acute SIV infection had no effect on the percentage of FJC+ cells throughout the brain. (B) The average cell intensity of FJC staining relative to DAPI staining is elevated in the putamen, thalamus, and hippocampus following acute SIV infection. (C) Average cell intensity of FJC correlated with markers of biological aging in the putamen, thalamus, hippocampus, and cerebellum. For A-B, all data points are presented and mean +/− SD are plotted for each group. Each brain region was analyzed independently using unpaired t-tests. For C, independent Pearson’s correlation coefficients were calculated using two-tailed p values between all of the markers for each brain region. Data for FJC intensity for all brain regions was then compiled into a single table. * p < 0.05, ** p < 0.01, *** p < 0.001.

The increase in FJC intensity also correlated with several markers of biological aging (Figure 5C). In the putamen, increased FJC intensity correlated with increases in the %p16+ cells (r = 0.861, p = 0.028), %p16+ astrocytes (r = 0.870, p = 0.024), p16 intensity (r = 0.862, p = 0.027), SIRT1 intensity (r = 0.897, p = 0.015), and GFAP intensity (r = 0.889, p = 0.018). In the thalamus, increases in the %p16+ cells (r = 0.924, p = 0.008) and %p16+ astrocytes (r = 0.923, p = 0.009) correlated with increased FJC intensity. In the hippocampus we saw both positive and negative correlations with FJC intensity, with the %p16+ cells (r = 0.896, p = 0.040), p16 intensity (r = 0.990, p = 0.001), and GFAP intensity (r = 0.985, p = 0.002) increasing with increased FJC intensity, while %SIRT1+ cells (r = −0.930, p = 0.022) and %SIRT1+ astrocytes (r = −0.935, p = 0.020) decreased with increased FJC intensity. Finally, GFAP intensity in the cerebellum increased with increasing FJC intensity (r = 0.940, p = 0.005).

## Discussion

In this study, we examined six brain regions in RMs that are implicated in HAND. We found significant alterations of the aging markers p16 and SIRT1 with acute SIV infection, combined with neurodegeneration in select brain regions. The astrocyte marker GFAP increased with infection in most of the brain regions examined (Figure 2). Expression of p16 was significantly elevated with SIV infection across five of the six brain regions for both the total cell population and astrocytes specifically (Figure 3). Conversely, the percentage of SIRT1+ cells decreased in the hippocampus with acute infection, suggesting a more region- and disease-specific response for SIRT1 expression (Figure 4). Additionally, neurodegeneration significantly increased in acutely-infected animals relative to naïve animals across several brain regions (Figure 5). Furthermore, the expression of p16 and SIRT1 correlated with neurodegeneration in both the total cell population and in astrocytes across several brain regions (Figure 5). Together, these data provide critical new insights into the effects of acute viral infection on aging-related changes throughout the brain.

The intensity of GFAP expression (rather than the percentage of GFAP+ cells) is often considered a marker of astrogliosis and increases with eugeric aging in many species from rodents through Non-Human Primates (NHPs) to humans^25–27^. In this study, both the intensity and the percentage of GFAP+ cells increased during acute SIV infection in the caudate, putamen, and hippocampus. The increase in the percentage of GFAP+ cells could be due to the overall increase in GFAP intensity, as this could increase the number of cells that surpassed the threshold set to be considered GFAP+. When combined with p16 data, these results suggest that the caudate, putamen, and hippocampus may be highly sensitive to the aging effects of acute infection with SIV. As we^20^ and others^28,29^ have shown that there is an accelerated rate of aging following S/HIV infection that can be reduced with therapeutics^30^, including cART^20^, these studies emphasize the need for early cART intervention.

In the CNS, p16 increases with normal aging in the frontal lobe of both human^31^ and NHPs^24^, especially in astrocytes^24,32^. We and others have also described region-specific increased expression of p16 following exposure to S/HIV^6,20^. Within astrocytes, *in vitro* exposure to HIV proteins increases expression of p16^33,34^, indicating this may also be an acute phase process. However, astrocyte expression of p16 throughout the brain during the acute phase of a viral infection in NHPs had not been previously analyzed. Here, we demonstrate robust increases in astrocytic p16 during early SIV infection across four brain regions implicated in HAND. These increases are also seen in the total cell population, which warrants further investigation to determine if other cell types experience similar significant increases in p16 expression with SIV infection or if this is an astrocyte-driven response. Importantly, expression of p16 could be indicative of cellular senescence in these cells which would prevent them from performing critical homeostatic functions – including synaptic regulation and blood-brain barrier function – and lead to damaging effects in the CNS. However, as p16 expression alone is not sufficient to determine a cellular senescence phenotype these studies assessed the expression of additional molecular markers associated with senescence to better assess the status of these cells.

It is of interest that we observed a significant decrease in the levels of SIRT1 in the hippocampus, as SIRT1 acts largely by inhibiting several pathways associated with aging and is downregulated with age in several tissues^35^. As the HIV-1 Tat protein is known to directly interact with SIRT1 and inhibit its activity^36^, decreased expression combined with a decrease in activity could lead to a drastic acceleration in aging processes. Further, we observed neurodegeneration, again in the putamen, thalamus, and hippocampus – brain areas linked to degeneration^37^ and cognitive decline in HAND.

These studies have raised an intriguing question: Can acute events such as bacterial and viral infections that resolve relatively quickly produce accelerated aging phenotypes within the CNS? Our previous studies have shown that astrocytes remain activated following infection and clearance of several infections including Brucella, Chikungunya and Dengue^38–40^. Further, certain psychiatric conditions also have similar changes in glial activation that can be reversed with antidepressant medications^41,42^. While those studies were not set up to address the specific question of accelerated aging, it does open a potential new avenue for research, with new targets for such insults.

These findings may hold clinical relevance for PLWH, and potentially for people exposed to HIV, but who do not necessarily seroconvert or otherwise progress. Human imaging studies on brain aging in PLWH have demonstrated premature, accentuated, or accelerated aging throughout regions of the brain implicated by the cognitive and motor deficits seen in HAND, including the frontal cortex, caudate, putamen, hippocampus, thalamus, and cerebellum^1,2,22,43–52^. Importantly, cognitive decline also shows premature onset or accelerated progression in PLWH^3,53^ and SIV-infected RMs^10,11,16,19,21,54–57^.

Overall, this study demonstrates for the first time that multiple markers of aging are significantly elevated, especially in astrocytes, during acute SIV infection across several brain regions implicated in HAND. Additionally, these alterations are correlated with neurodegeneration in the putamen, thalamus, hippocampus, and cerebellum. Given the importance of these brain regions for learning/memory and executive functioning, two cognitive domains that have been shown to be significantly more impaired in PLWH in the cART era than pre-cART^7,58^, our results may have critical implications in the treatment or prevention of HAND. Longitudinal work is needed to determine the direct effects of these aging markers on the progression of neurodegeneration and development of HAND and the underlying mechanisms at play.

## Author Contributions

MDH selected tissues from the extensive archive, designed the study, performed immunofluorescent staining, analyzed data, and wrote the manuscript.

CCM performed Iba1 staining and drafted parts of the manuscript.

ARVZ assisted with immunohistochemistry and drafted parts of the manuscript.

AAS performed histological analyses to ensure unbiased selection of tissues, and drafted parts of the manuscript.

AGM oversaw all aspects of the study.

## Acknowledgements

We would like to thank Dr. Robert Blair for training on the use of HALO® image analysis software, the staff of the Anatomic Pathology Core RRID: SCR_024606, Clinical Pathology Core RRID: SCR_024609, Confocal Microscopy and Molecular Pathology Core RRID: SCR_024613, and Virus Characterization, Isolation, Production and Sequencing Core RRID:SCR_024679. This work could not happen without the dedicated staff and veterinarians of the Division of Veterinary Medicine at TNBRC RRID: SCR_008167.

